# Ratio Percentile Deviation (RPD): A nonparametric, compositionally robust method for measuring the divergence of a microbial sample from a reference dataset

**DOI:** 10.64898/2026.05.27.728224

**Authors:** Cristina M. Herren

## Abstract

Comparing a microbial sample against a reference dataset is essential for many workflows, such as quantifying change in a microbial community after experimental perturbation or evaluating whether a human microbiome sample falls within the range of healthy subjects. Here I present a new method called Ratio Percentile Deviation (RPD) for quantifying the divergence of a microbial sample from a set of reference samples. The RPD method compares features of the test samples to the empirical distribution of features in the reference samples, making no assumption about the underlying distribution of the data. The features used for comparison are ratios between taxa in the same sample, which are invariant between count and relative abundance data; therefore, RPD is appropriate for compositional read-based data. Applying the RPD method to several large microbial datasets shows that it outperforms approaches that compare test samples to a composite reference value (e.g. the centroid of the reference dataset). In case-control analyses using the American Gut Project and Ghana Breast Health datasets, a single RPD predictor discriminated individuals with microbiome dysfunction from healthy controls with AUCs of 0.74-0.79; DeLong tests confirmed RPD’s AUC was significantly higher than the best-performing comparator in all three analyses. I further show that RPD can measure compositional variation within a dataset by applying the metric to the long-term Lake Mendota microbial timeseries. The superior predictive power of RPD versus existing approaches across these varied datasets suggest that this method could be a useful new tool for comparing microbial samples to a reference across multiple study systems. The RPD method is available as an R function in the following github repository: https://github.com/cherren8/RPD

## Introduction

A consistent challenge in the analysis of microbial data is compressing large datasets into a small number of meaningful values to use in statistical models (Ramette 2007, Armstrong et al. 2022, Fuschi et al. 2025). Microbial community composition samples are generally in the form of vectors, with hundreds to thousands of observations per sample or community. The problem of comparing two communities in an ecologically meaningful way has several widely accepted solutions, including Bray-Curtis dissimilarity (for compositional data [Bray and Curtis, 1957]), Unifrac distance (for incorporating phylogenetic data [Lozupone and Knight, 2005]), and Jaccard distance (for presence/absence data [Jaccard, 1912]). These metrics provide a single measurement of the divergence of the two vectors. However, a more difficult problem is quantifying how different one community or vector is from a set of reference communities. Despite the widespread need to query microbial samples against a database, there is less consensus on methodological approaches for this problem (Schloss 2026, Kumar at al. 2026).

Comparing a test sample (or samples) to a reference occurs in a number of important applications, including: comparing the divergence of the microbiome of individuals with microbiome dysfunction to the microbiome of a healthy population; quantifying whether one of many experimental treatments shifts community composition further from baseline; or measuring the deviation over time of microbial community composition in response to environmental stressors. A commonly used approach is to use some form of averaging on the reference set, and then use a distance or dissimilarity metric from the test sample to the composite value from the reference set (Anderson et al. 2006, AlShawaqfeh et al. 2017, France et al. 2020, Schloss 2026). An example would be to take Bray-Curtis dissimilarity of each test sample to a theoretical sample that is the mean of all the reference samples (e.g. Widder et al. 2022, Frøslev et al. 2023, Hoy et al. 2026). However, this approach has the critical drawback that the centroid community does not actually exist in nature; the centroid community is created primarily because it enables the use of a distance or dissimilarity metric. There are many cases where this approach of averaging across the reference is suboptimal. For example, in temperate biomes, there is seasonality in the microbial community (Kara et al. 2013, Landesman et al. 2019). Averaging the composition of a year’s worth of samples will yield a community that averages across this seasonality, yielding a centroid community that would not naturally occur. Taking the distance to this centroid would contain little ecological information.

This paper presents a new metric for quantifying the deviation of a microbial sample from a reference dataset. It uses the reference dataset to build empirical distributions of features in the data, and then compares the test sample to these empirical distributions (Fig. 1). Specifically, the features of the reference data considered here are the ratios between taxa within the same sample (e.g. the ratio of taxon 1 to taxon 2, or taxon 4 to taxon 65). Ratios within samples are identical between absolute value or relative abundance data, meaning that they are appropriate for analyzing compositional microbiome data (Gloor et al. 2017). Additionally, a long history of ecological modeling and theory suggests that ecological dynamics should be governed by the ratios of taxa in addition to their absolute abundances (Rosenzweig 1971, Holt and Pickering, 1985, Arditi and Ginzburg, 1989). Several empirical microbiome studies have found that altered ratio of bacterial taxa in the human gut are associated with disorders such as anorexia nervosa (Helal et al. 2024), autism spectrum disorder (Kadiyska et al. 2025), and multiple sclerosis (Ghimire et al. 2025). Thus, there is both empirical and theoretical underpinning to support that the ratios of taxa within the same sample are informative measurements of a community. The method presented here calculates the pairwise ratios in all the reference communities, and then finds the percentile in this distribution where the test sample falls. It then calculates the deviation from the median in these percentiles, with higher deviation meaning that the sample has more extreme values in comparison to the reference set (Fig. 1). As such, the method is named Ratio Percentile Deviation (RPD).

**Figure 1:**
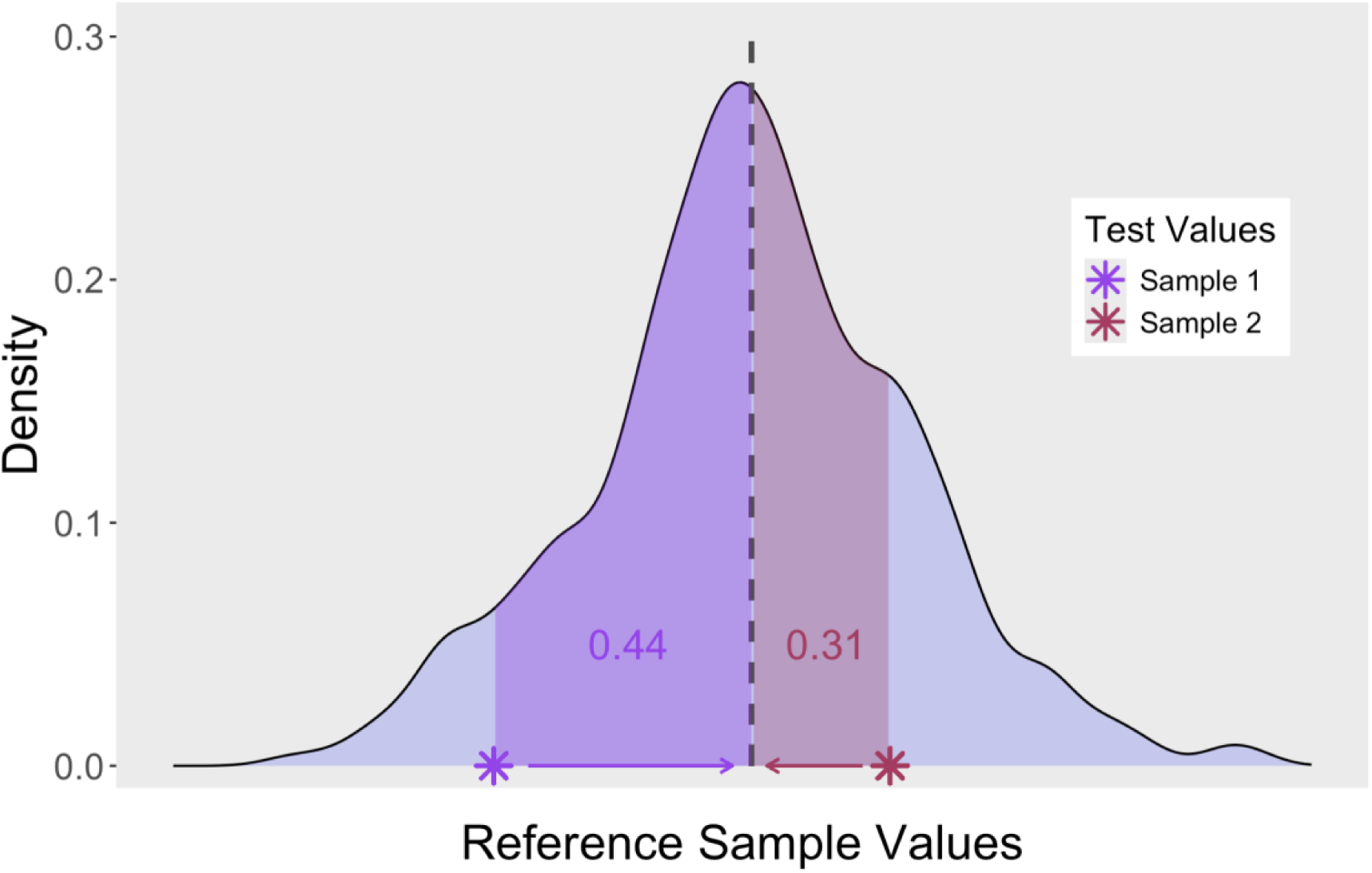
Comparison of test values to empirical distribution of sample values. The blue density plot shows a distribution of empirical values calculated from reference data. Two test values are overlaid onto the plot, which fall at the 6th percentile (Sample 1, purple) and the 81st percentile (Sample 2, red). The RPD method scores how far the test samples are from the reference set by calculating the fraction of the reference distribution that falls between the sample value and the median (50th percentile, dashed grey line). The shaded area corresponds to these scores (0.44 for Sample 1, 0.31 for Sample 2).

This paper presents the new RPD method and uses it in four applications across three large microbiome studies. Three of the applications use the method in a biomedical context, where RPD is used as a measurement of the difference between samples from individuals with microbiome dysfunction and samples from healthy references. In the final analysis, RPD is used to evaluate drivers of compositional variability in an environmental microbiome dataset. In this case, I examined whether RPD responds to known environmental forcing variables. Together, these four analyses across different ecosystems demonstrate the utility of RPD in analyses that compare microbial samples to a reference dataset.

## Methods and Results

### Overview

The RPD method takes inputs of a reference dataset and a set of test samples. It returns one score for each test sample quantifying how much the sample deviates from the reference dataset. Thus, the method is compatible as a predictor or outcome in a variety of subsequent statistical analyses. RPD scores are determined by comparing the ratios of taxon abundances in the test samples to the distribution of taxon abundance ratios in the reference samples. A higher sample RPD score indicates that taxon abundance ratios are more unusual or extreme in comparison to the reference dataset.

### Algorithm

Begin by separating the data into a reference set and a test set. The samples should contain the same taxa, and therefore have the same number of columns. The algorithm proceeds through the same steps for each taxon:

1. For a given focal taxon, divide all other abundances in each sample by the focal taxon’s abundance. This converts the data to ratios. Perform this calculation for both the test set and the reference set.
2. For each ratio in the test set, calculate the percentile where it falls in the corresponding distribution of reference ratios. Find this percentile value by counting the number of reference ratios that are smaller than the test ratio and dividing by the total number of reference ratios.
3. Calculate the absolute deviation from the median percentile (0.5) for all ratio percentiles in each sample.
4. Take the average of the absolute deviations. This is the taxon-level RPD score.

Repeat these steps with each taxon as the focal taxon. This will produce a vector of taxon-level ratio percentile deviation values for every sample. Taking the mean of these taxon-level RPDs yields the sample-level RPD. By default, the calc.rpd() R function returns a vector of RPD scores corresponding to the input test samples. An optional argument also returns taxon-level RPD scores for each sample.

A higher RPD indicates that taxa have more variability in their taxon abundance ratios. Larger RPD values also occur when percentiles are more extreme, meaning they are close to 0 or 1. A percentile value of 0 occurs when the test ratio is smaller than all the reference ratios, and a percentile value of 1 occurs when the test ratio is larger than all the reference ratios. The maximum RPD score of 0.5 occurs when all ratio percentiles are 0 or 1.

### Algorithm Parameters and Zero Handling

The large number of rare taxa in microbial datasets means that there must be criteria for excluding sufficiently uncommon taxa from the analysis (Herren 2019, Jia et al. 2022, Nearing et al. 2022). Thus, there is a reference persistence parameter that specifies what fraction of reference samples a taxon must occur in to be included in the analysis. Depending on the heterogeneity of the dataset and the depth of sequencing, this value generally ranges from 0.5 to 0.9. This parameter is necessary because is important for taxa to be present in enough samples to have a representative distribution of ratios in the reference set.

Because of the sparsity of microbial datasets, decisions about how to handle zero values can have a strong effect on the outcome of an analysis. The RPD algorithm only assigns taxon-level RPD values to taxa that have non-zero abundance in a sample, because a focal taxon abundance of zero creates undefined (infinite) ratios.

However, zero values are retained when the numerator of a ratio is zero. This occurs when there are zeroes in the non-focal taxon abundance (Fig. 2). A zero value in the numerator leads to a ratio of 0, which subsequently leads to a percentile of 0. Thus, a zero ratio necessarily leads to the largest possible deviation from the median, at 0.5. As fewer taxa overlap between the reference and test datasets, more ratios will have numerators of 0 and deviations of 0.5., and the sample-level RPD value will trend towards 0.5. As such, test samples with no taxa shared with the reference set receive an RPD value of 0.5.

**Figure 2:**
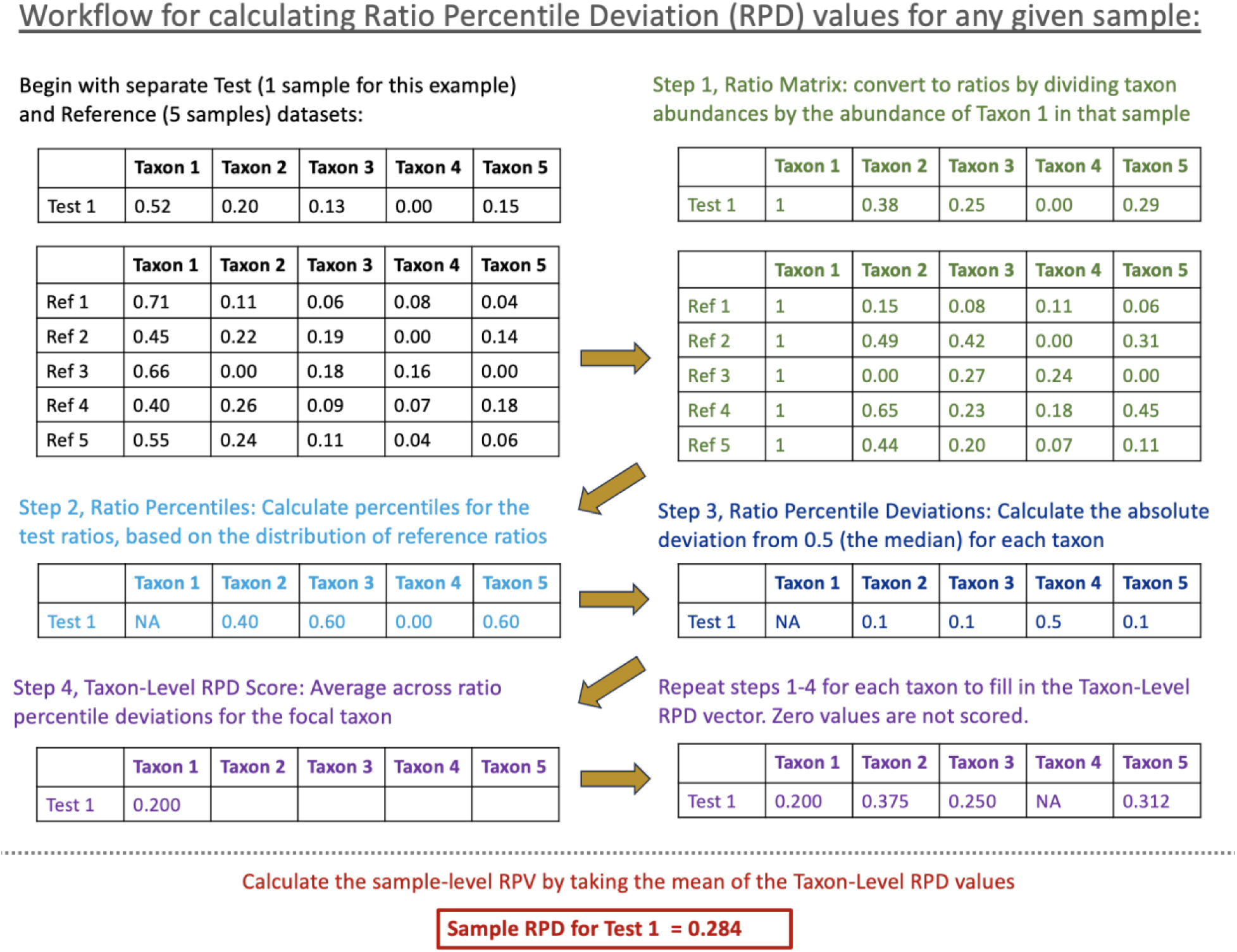
A schematic of the RPD algorithm. This figure shows the steps involved in calculating an RPD score for one test sample (Test 1) using a reference dataset of 5 samples (Ref 1-5). Steps correspond to those detailed in the *Algorithm* section.

### RPD for Variation Within a Dataset

To measure variation within a single dataset, RPD values can be calculated using the same samples as both the reference and test inputs. When the input and test datasets are identical, the RPD algorithm leaves out the focal sample when calculating the percentile in the reference set to avoid self referencing.

### Datasets Analyzed

I conducted four analyses using three large microbial datasets of different origins. Three of the four analyses look at the utility of RPD as a predictor of microbiome dysfunction/dysbiosis. For these analyses, I used the American Gut Project (AGP) dataset (McDonald et al. 2018) and the Ghana Breast Health (GBH) study (Byrd et al. 2021). The AGP dataset is a crowdsourced collection of human-associated samples (primarily stool) with corresponding metadata on a variety of lifestyle variables. The GBH study collected stool samples from women in Ghana at the time of breast biopsy, and found that individuals with breast disease had reduced diversity and differential abundance of microbial taxa associated with immune function.

These datasets were selected based on the following criteria: access to the same data analyzed in previous publications, inclusion of a large control population, large sample size, and variety of disorders studied. Data analysis scripts and abundance tables used for analysis can be found in supplemental material for all analyses.

In the three datasets evaluating whether RPD could predict microbiome dysbiosis, I divided each dataset into reference and test sets. The reference set for the RPD method was comprised of healthy individuals. The test set was comprised of control and case samples, where controls were additional healthy subjects and cases were individuals with a history of microbiome dysfunction (either a recent or present disorder).

I conducted two separate analyses with the AGP dataset: one with cases defined as individuals diagnosed with a *C. difficile* infection, and one with cases defined as individuals who used antibiotics within the last month. Both these scenarios are well documented as corresponding to microbiome dysbiosis (Antharam et al. 2013, Vasilescu et al. 2022, Fishbein et al. 2023, Cusumano et al. 2025). For both these analyses, a healthy reference dataset was established using adult individuals from the United States who did not report gastrointestinal disorders, use of antibiotics, or cancer. To reduce dataset heterogeneity and for quality control, the data were additionally subsetted to individuals within the United States who provided a height and weight and samples with a minimum of 2000 reads. The version of data downloaded was the deblurred data trimmed to 125 nucleotides with post-collection blooms removed. Taxa present in fewer than 5 samples were removed from the dataset. These data correspond to the dataset analyzed by McDonald et al. 2018.

For the GBH study, individuals were divided into case and control by whether they had a diagnosis of breast disease. Data were subsetted to samples with greater than 10000 reads. The version of data downloaded used OTU picking and a trimmed length of 100 nucleotides. The discrepancy in read cutoff between this and the AGP dataset was driven by a difference in average read sequencing between this dataset and the older AGP dataset. These data correspond to the dataset analyzed by Byrd et al. 2021.

For the final analysis, I selected an environmental microbiome dataset to test the RPD method in a non-human system. The Lake Mendota microbiome dataset is one of the longest and best-documented time series of an environmental microbiome, including hundreds of samples with matching dissolved nutrient data (Rohwer et al. 2023, Rohwer et al. 2025). In this analysis, I asked whether community composition deviated in response to nutrient variability. I used RPD as a metric of “deviation” in community composition, and measurements of dissolved nitrogen and dissolved phosphorus as the nutrient data. I subsetted the data to the well-sampled months of May through October for the best-sampled years 2013-2018. These data correspond to the dataset analyzed in Rohwer et al. 2023.

### Benchmarking and Sensitivity Analysis

The results of analyses involving RPD are compared to equivalent analyses carried out with Bray-Curtis Dissimilarity (BCD) to the reference centroid in place of RPD. For further benchmarking, the supplementary materials present comparisons to the same analyses carried out using rank shift analysis (Collins et al. 2008), robust Aitchison distance to the reference centroid (Aitchison 1986, Martino et al. 2019), median Bray-Curtis Dissimilarity to all references (Lloyd-Price et al. 2019), and Euclidean distance. I also benchmarked against Bray-Curtis Dissimilarity and robust Aitchison distance calculated on the subset of taxa retained for the RPD analysis. The primary method of evaluation of the various predictors was the area under the curve (AUC) for the Receiver-Operator Curve (ROC) when classifying case versus control status using each predictor (Hanley and McNeil, 1982). Bray-Curtis Dissimilarity to the reference centroid had the best performance out of the benchmarking comparisons, which is why it is presented here.

The supplementary materials also provide the following sensitivity analyses: effect of repeated random sorting of healthy individuals into reference and control samples; effect of the reference persistence parameter; effect of the size (number of samples) of the reference dataset; effect of model optimism by validation on hold-out data. None of these effects qualitatively changed the results presented in the main text.

### RPD Values and Relationship to Diversity and BCD

There were 699 individuals in the American Gut Project dataset who met the criteria for control samples. Of these, a random sample of 500 were taken as the reference group. The remaining 199 were used as the control samples in the test set.

For the AGP antibiotics analysis, there were 321 individuals who reported taking antibiotics within the last month, which comprised the case portion of the test dataset. I calculated RPD values using the set of 500 healthy individuals as the reference data, and the set of 199 controls and the 321 individuals with recent antibiotic use as the test data. The reference persistence threshold used was 0.7. RPD values for control individuals had a mean of 0.273 with a standard deviation of 0.0528 (Fig. 3). The mean RPD value for individuals with antibiotic use within the last month was 0.331 with a standard deviation of 0.0720.

**Figure 3:**
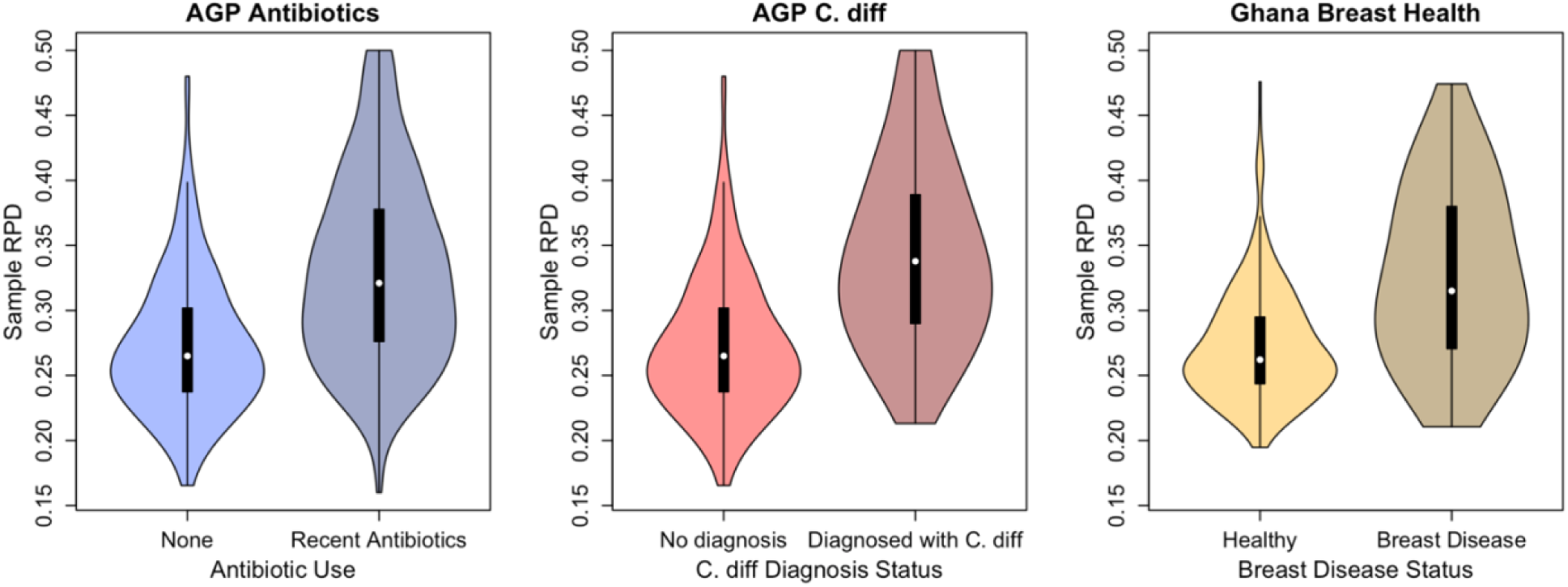
Violin plots of RPD values of cases and controls for the three datasets: AGP antibiotics, AGP *C. difficile*, and GBH. For all three datasets, the mean RPD for controls (left in each panel) are substantially lower than for cases (right in each panel). Colors match Figure 4, where darker shades in each panel indicate cases.

For the AGP *C. difficile* analysis, I used the same reference and control datasets, but used the 76 individuals with a diagnosed *C. difficile* infection as the case data. The reference persistence threshold used was again 0.7, meaning the control RPD values were identical to those in the AGP antibiotics analysis. Individuals with a *C. difficile* diagnosis had a mean RPD of 0.343 (standard deviation: 0.0717).

Finally, for the GBH study, a random sample of 100 healthy individuals were used as a reference, and the test set was comprised of the remaining 310 healthy individuals and the 509 individuals with breast disease. The reference set selected was smaller here due to the more limited number of healthy controls. However, the greater homogeneity of the dataset compensated for the smaller size of the reference dataset (see SOM). The reference persistence threshold used was 0.7. The mean RPD values for controls was 0.273 (standard deviation: 0.0430) and for cases with breast disease was 0.326 (standard deviation: 0.0651).

Across all three datasets, RPD was inversely related to alpha diversity and positively related to Bray-Curtis Dissimilarity to the reference centroid (Fig. 4). Although correlations were clear and consistent, the strength of the relationship with diversity was moderate, as correlation values with diversity were -0.576 (AGP antibiotics), -0.541 (AGP *C. difficile*), and -0.496 (GBH). The relationship between BCD and RPD was stronger, with correlation values of 0.800 (AGP antibiotics), 0.780 (AGP *C. difficile*) and 0.848 (GBH).

**Figure 4:**
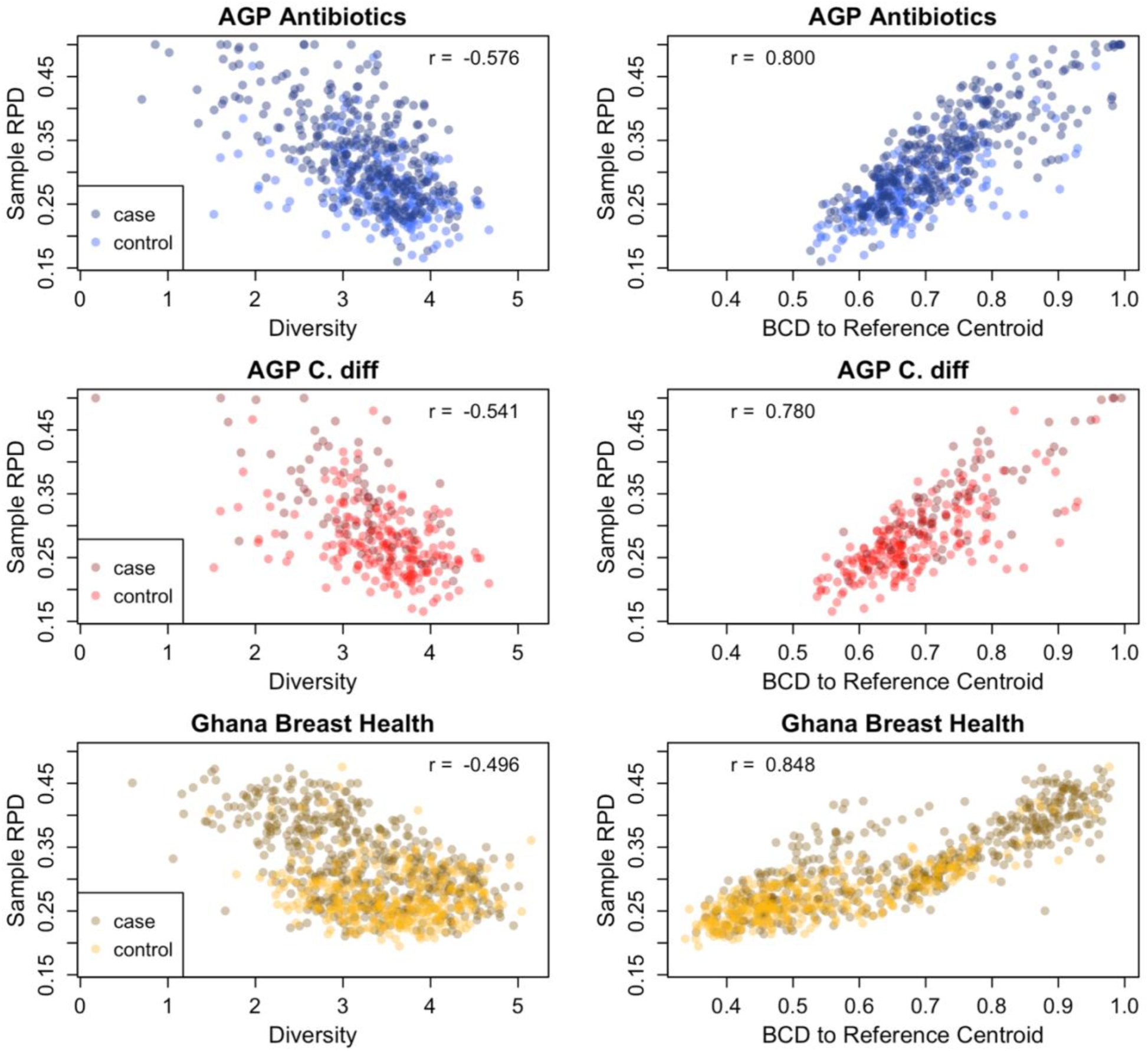
Correlations between RPD and standard microbiome metrics (alpha diversity and BCD to reference dataset centroid). In all three datasets, there is a moderate negative correlation between RPD and diversity, where less diverse samples are associated with higher RPD. The relationship between BCD to the reference centroid and RPD is stronger, where samples with higher BCD also have higher RPD. Darker colors within each plot indicate cases, whereas lighter shades within each plot indicate control samples.

### RPD as a Predictor of Microbiome Dysbiosis

To investigate the utility of RPD as a predictor of case vs. control status, I created three logistic regression models (one for AGP antibiotics, one for AGP *C. difficile*, and one for Ghana Breast Health) using 0 for controls and 1 for cases. The predictor was the RPD value for each sample. I then obtained the AUC for the associated ROC plot from the fitted models. I compared the model significance and AUC to alternative models fit using sample BCD to the reference centroid as the predictor.

RPD was a better predictor of case vs. control status than BCD and was highly significant across all three analyses (Fig. 5). For the AGP antibiotics study, RPD was significant in the logistic regression at p = 7.46-17 and had an AUC of 0.744. As a comparison, BCD as a predictor was significant at p = 2.46e-8 and had an AUC of 0.651. For the AGP *C. difficile* study, RPD as a predictor was significant at p = 1.51-11 and had an AUC of 0.789, whereas BCD as a predictor was significant at p = 2.03e-7 and had an AUC of 0.718. Finally, for the GBH study, RPD as a predictor was significant at p = 9.95e-27 with an AUC of 0.745, while BCD was significant at p = 1.91e-21 with an AUC of 0.706.

**Figure 5:**
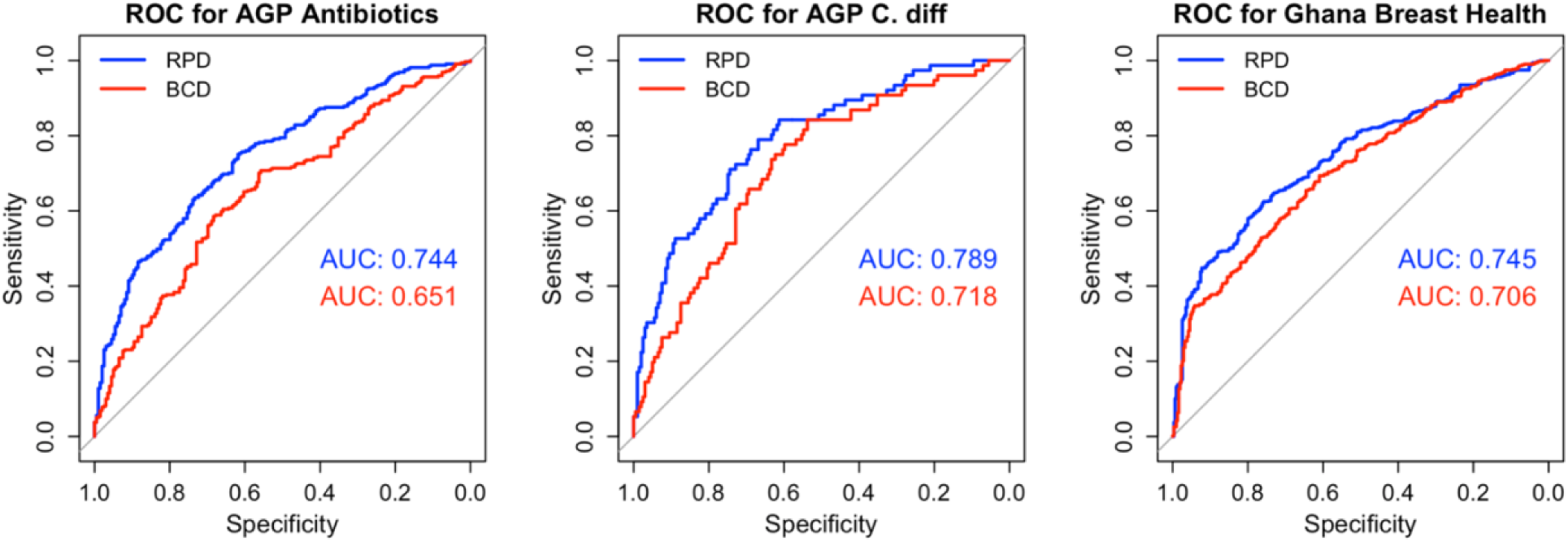
ROC plots and associated AUC values for logistic regressions modeling case vs. control status as a function of either BCD or RPD. For all three datasets, RPD is a significantly better predictor of case vs. control status, as measured by AUC and associated DeLong tests. The greatest difference in AUC values between models is in the AGP antibiotics dataset, with a difference of 0.093 (AGP *C. difficile*: 0.071; GBH: 0.039).

I implemented DeLong’s test (DeLong et al. 1988) to statistically compare the AUC values produced by the RPD and BCD analyses. This test accounts for the fact that the predictors are fit to the same data, and therefore have correlated predictions. In all three cases, the AUC for RPD as a predictor was significantly higher than the AUC for BCD as a predictor (p = 6.79e-8 for AGP antibiotics; p = 0.0034 for AGP *C. difficile*; p = 0.00064 for GBH).

High RPD values were strongly related to a higher risk of microbiome dysbiosis. For the AGP antibiotics analysis, the relative risk was 1.77 for samples in the top half of RPD scores. For the AGP *C. difficile* dataset, this relative risk was 4.40, and for GBH it was 1.79.

### Drivers of Lake Mendota Microbial Compositional Variability

The final analysis evaluated the sensitivity of RPD as a method for measuring compositional variability within a dataset. I used the Long-Term Ecological Research Microbial Observatory 16S data from Lake Mendota with paired measurements of dissolved N and dissolved P as environmental predictors. I selected this dataset because of the known importance of dissolved nutrients as drivers of freshwater microbial community composition (Chrost et al. 2009, Knelman et al. 2014), including in Lake Mendota specifically (Newton and McMahon 2011). Furthermore, dissolved N and P values in Lake Mendota vary considerably, both over a season and between years (Fig. 6, Brock 2012, Soranno et al. 1997). Although there is seasonality in microbial community assembly in Lake Mendota, there is also year-to-year variation in composition, particularly during the transition from spring to summer dynamics (Fig. 6). I asked whether compositional variability, measured by RPD, was related to nutrient variability.

**Figure 6:**
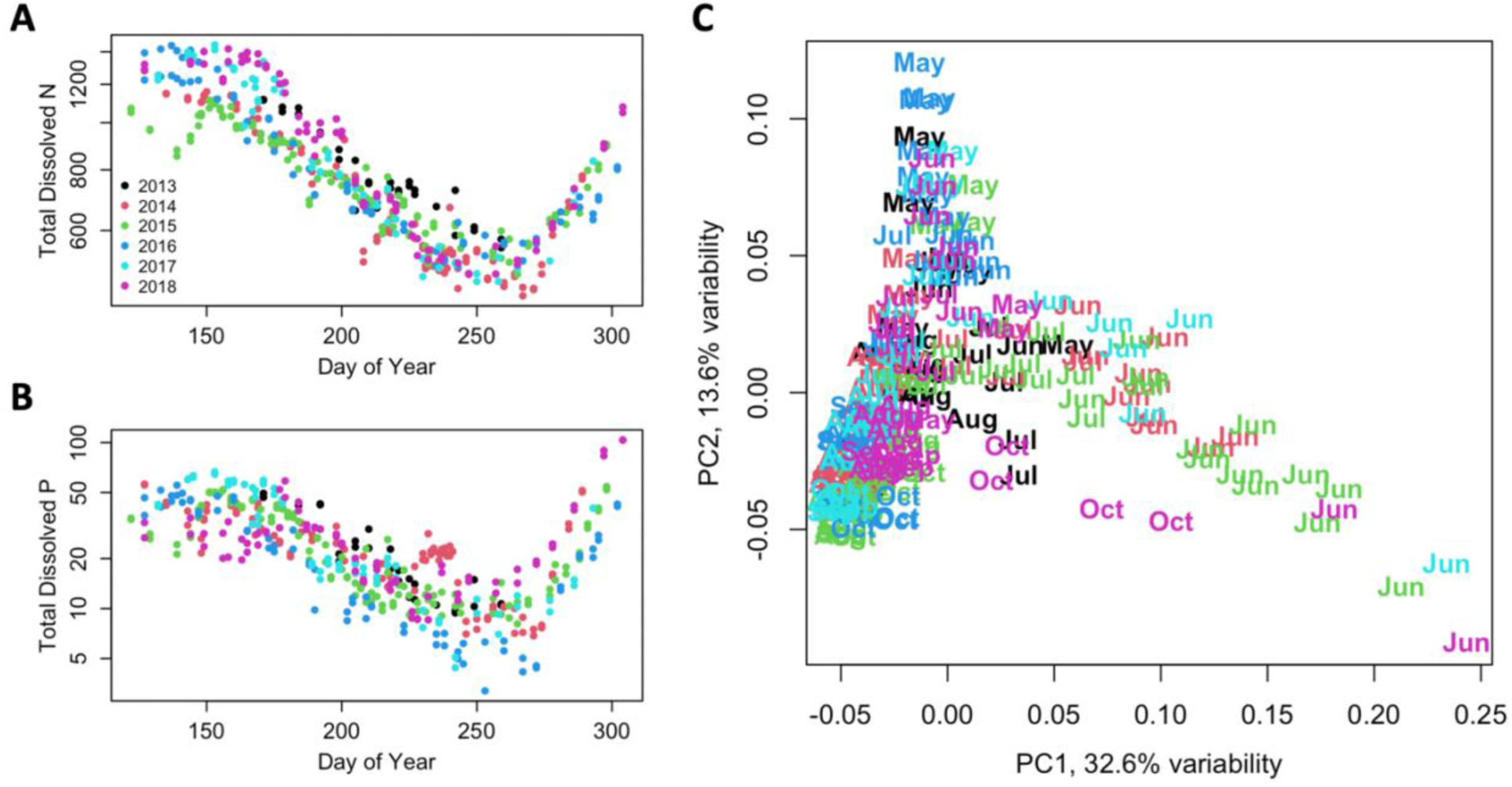
Seasonal and inter-annual dynamics of dissolved nutrients (panels A and B) and microbial community composition (panel C) in Lake Mendota for 2013-2018. There is both within-year and between-year variation in dissolved nutrient availability, with a greater than 3-fold difference in dissolved nitrogen values (panel A) and a greater than 10-fold difference in dissolved phosphorus values (panel B) across the dataset. Microbial community composition is captured in a PCA with the first and second axes representing a total of 46.2% of dataset variation (panel C). The first axis is associated with variation in community composition in the early summer, mainly June, while the second axis is associated with variation in late spring, mainly May. Colors represent different years.

For this analysis, the reference data and the test data were the same, because the goal was to evaluate how much each sample deviated from the remainder of the dataset. There were 232 samples that met the inclusion criteria of being from the months of May to October in the years 2013-2018. For these same dates, there were matching dissolved N and dissolved P values for 222 samples. Nutrient samples were taken in duplicate, which was used for quality control of the data. I calculated the standard deviation of the log-transformed paired values for dissolved N and dissolved P. There were clear high outliers for 5 dissolved N pairs and 3 dissolved P pairs. Of these pairs, there was always one sample that fell outside the seasonal trend (visible in Fig. 6). The anomalous point was removed for these 8 pairs. For all other pairs, values were averaged to produce one dissolved N and one dissolved P value per sample date.

I asked whether more extreme dissolved nutrient concentrations corresponded to more unusual community composition, where unusual community composition was measured by RPD. The persistence threshold used in this dataset was 0.85. For the nutrient data predictors, I first calculated the z-scores of the log-transformed dissolved nutrient concentrations. This gave a distribution of how far the sample deviated from the mean dissolved nutrient value over this time period. Scores further from 0 (either positive or negative) indicated greater deviation from the average. Thus, my final metric of nutrient anomaly was the absolute value of these z-scores. On this scale, zero is the most typical nutrient value, with higher values indicating greater deviation from average.

I regressed RPD against the z-score absolute values for dissolved N and dissolved P. I performed the same analysis instead using BCD to the dataset centroid as the measurement of compositional deviation. The analysis using RPD as a response variable had a much stronger relationship with nutrient deviation values, with significance of both the N and P variables (p < 0.001 for N, p = 0.0014 for P) and an adjusted R^2^ value of 0.33. In the parallel analysis instead using BCD as the outcome, the adjusted R^2^ value was 0.17 and only the N deviation variable was significant (p < 0.001 for N, p = 0.057 for P).

### Software, Packages, and Data

Analyses were carried out in the R Programming Language, version 4.5.1. Packages used included ggplot2 (version 4.0.3) for generating Figure 1; vegan (version 2.7-3) for calculation of BCD and robust Aitchison values, as well as diversity values; vioplot (version 0.5.1) for violin plots of RPD values; pROC (version 1.19.0.1) for ROC plots, AUC values, and DeLong tests; lubridate (version 1.9.5) for handling date conversions; and limony (version 0.0.0.9000) for accessing Lake Mendota data.

Data for the AGP and GBH studies were accessed through the Qiita portal. Lake Mendota 16S data were accessed through the limony package. Lake Mendota nutrient data were accessed through the Environmental Data Initiative online portal.

## Discussion

### Interpretation of Results

Across all four datasets, RPD enabled better statistical modeling of microbiome dynamics than Bray-Curtis dissimilarity. In the three cases of RPD as a predictor of microbiome dysfunction, the areas under the curve in the ROC analyses were between 0.74 and 0.79; this ability to differentiate cases from controls is notable given that only one predictor was included in the model. Furthermore, the improvement in area under the curve from BCD to RPD was more than 0.05 for two of the datasets, indicating a meaningful improvement. For the final dataset, although the improvement was more modest at 0.039, this change in model fit was still strongly significant. Supplementary analyses demonstrate that model optimism is below 0.005, meaning that average RPD performance was almost identical on held out data. Thus, RPD has strong potential as a predictor in microbiome classifier models, or as a screening criterion to identify individuals with anomalous microbiome composition.

In the Lake Mendota dataset, RPD responded more strongly than BCD to known drivers of compositional change. This suggests that RPD is a more sensitive measurement of compositional variability in this system. Furthermore, RPD showed a significant relationship with both deviation in dissolved N and deviation in dissolved P, whereas BCD showed only a significant relationship with dissolved P. Thus, the heightened sensitivity of the RPD metric enabled the detection of a secondary relationship that was not captured when measuring compositional variation with BCD. The relationship between RPD and BCD was strong and positive in all datasets, with correlations near 0.8 in all cases; this is encouraging, given that BCD to the centroid is a widely applicable and reasonably well-performing approach (Schloss 2026). One possible reason that RPD may have outperformed BCD as a predictor is because of the seasonality in the data. Averaging across the seasonal compositional variation yields a centroid that is not close to any of the individual data points, visible by the fact that the lowest BCD to centroid values are around 0.4 (Fig. 7). Therefore, RPD may be a superior predictor to centroid-based methods in datasets where there are multiple “normal” or “healthy” states.

**Figure 7:**
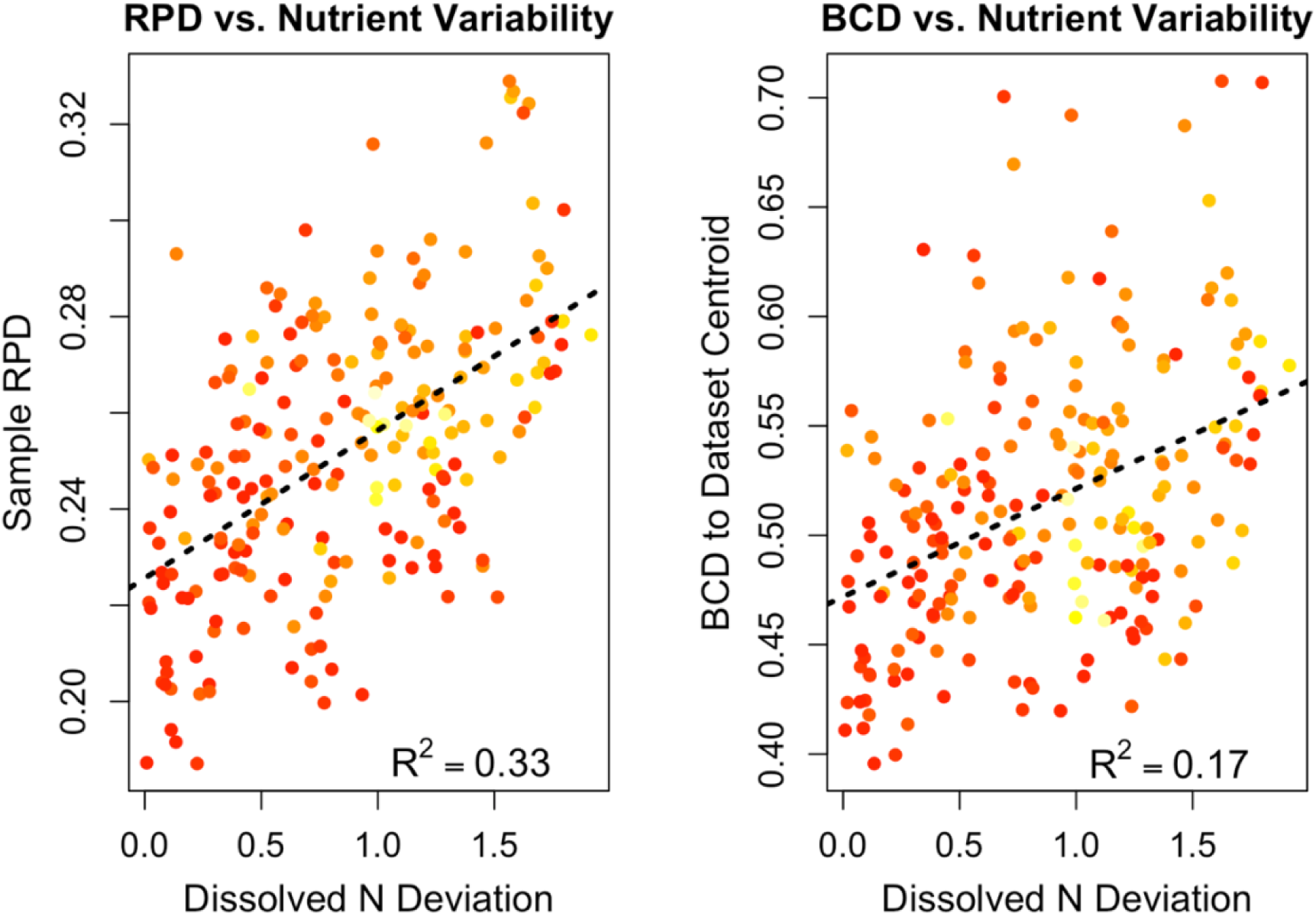
Relationship between metrics of compositional variability (RPD and BCD) and dissolved nutrient anomaly. Multiple regressions were carried out between each of the two metrics of compositional variability and paired dissolved N and P deviation values. The linear relationship between the metrics and dissolved N values are shown. Colors indicate associated dissolved P deviation values, where darker colors (red) indicate lower deviation and lighter colors (yellow) indicate higher deviation. The multiple regression adjusted R^2^ values are given.

### Comparison to Existing Methods and Advantages of RPD

In addition to the differences with BCD that were highlighted already, there are further ways in which RPD differs from centroid-based approaches to quantifying compositional variation. The most prominent difference in the RPD method is that it creates an empirical distribution from reference data. This avoids any assumptions about the underlying distribution of the populations or what distance measurement is most meaningful. Furthermore, distance or dissimilarity-based methods can change substantially when implemented on absolute abundance vs. relative abundance data. The RPD method performs identically, because the ratios of taxa are unchanged between absolute and relative abundance versions of the data. Given that the impact of absolute abundance change on microbial community analysis is generally unknown and unaccounted for, this is a strong point in favor of the RPD method. Finally, the RPD method is more robust to blooming taxa than distance-based metrics or BCD. A single high-abundance taxon can change distances or BCD substantially; in RPD, the ratios between all non-blooming taxa remain unchanged, leading to greater stability of the measurement in the presence of high-variance taxa.

The RPD method is easy to implement and integrates well with downstream statistical analyses. The function to calculate RPD uses only the reference dataset, test dataset, and a single parameter for the reference persistence threshold. It generates one value for each sample, which retains the appropriate sample size for later statistical analysis. Examples can be found in analyses of microbial datasets where multiple datapoints are generated from each sample, such as taking the BCD from the test sample to each of the reference samples. These approaches are not statistically valid, because they treat the resulting data points as independent when they are not. The RPD approach avoids this pitfall. Finally, the RPD method is fast, as RPD scores for the three datasets were calculated in 6.7 seconds (AGP antibiotics), 7.8 seconds (AGP *C. difficile*), and 100.4 seconds (GBH) using a laptop computer with 16GB RAM.

### Additional Applications

This new method has the potential for application to a number of important research questions across several domains of microbiology. As noted already, RPD might be used as a preliminary screening tool for disorders involving microbiome dysbiosis; individuals with an RPD above some threshold may be considered for follow-up testing. If a reference dataset were available for individuals with a specific disorder, a second calculation of RPD using the disorder dataset as a reference might improve classification performance. Additionally, RPD could be used in ecological studies of resistance and resilience to disturbance, where resistance could be measured as the change in RPD after disturbance and resilience is the rate of decline in RPD after disturbance. The RPD method could also be applied in experimental studies that investigate the effects of various treatments on microbial community composition. In this scenario, RPD scores could be compared across treatments, using the same baseline dataset as the reference. This would enable detection of differential effects of treatments on the degree of shift in the microbial community. There are also cases where RPD may be an outcome, instead of a predictor. For example, population variability is theorized to increase before critical transitions or ecological tipping points (Carpenter and Brock, 2006), and RPD could represent a new way to quantify this community-level variation. If asking the question of which taxa are driving the observed patterns in any of these analyses, the taxon-level scores could indicate which taxa have the most extreme ratios. Finally, RPD could be used on other forms of data besides OTU or ASV tables; this method could be implemented with other read-based datasets such as metagenomic gene coverage or RNA-seq data.

### Limitations and Considerations

The primary limitation of the RPD method is the need for a representative reference dataset that spans the state space of control communities. The following conditions will negatively impact the performance of RPD: systematic variation between the reference dataset and controls in the test set, smaller size of the reference set, non-representative sampling within the reference set. The appropriate size of the reference set depends on the homogeneity of samples and the depth of sequencing. Both these factors impact the number of taxa that are highly persistent, meaning they are detected in a large fraction of samples. For deeply sequenced and homogenous samples as might be obtained from an experimental study, it is possible that a reference set of 10-20 samples might suffice. However, as datasets become more heterogeneous, such as those from environmental samples or human-associated microbiota, it might be necessary to have 50-100 samples. The supplementary materials provide a sensitivity analysis on how the outcome of the AGP analyses change as reference size varies; the effect of reference size was relatively small when changing between a reference size of 20 and a reference size of 200. This is especially notable given that the AGP dataset is on the high end of dataset heterogeneity, being contributed by individuals across the United States with few barriers to inclusion. However, the effect of the reference size may become more important when there is less separation between cases and controls, because greater resolution may be necessary to differentiate samples.

The reference persistence threshold in the RPD function will likewise need adjustment based on the heterogeneity and depth of sequencing of the reference set. With greater heterogeneity, the persistence threshold must go down to retain a sufficient number of taxa. However, as sequencing becomes deeper, the mean persistence level in the dataset increases, meaning that the persistence parameter should be raised. In samples with very deep sequencing, it is likely that researchers will need to also subset their data based on a mean abundance threshold, as even very low abundance taxa might be detected in a majority of samples. In this case, taxa detected below some mean threshold should be removed before creating the reference and test datasets.

Finally, test samples that are extremely different from the reference set would all receive the same score of 0.5 if there are no taxa overlapping with the reference set. This may be the case if some test samples have extreme dysbiosis, such as sepsis (Miller et al. 2021). It would be a mistake to consistently remove these data from an analysis, because their lack of overlap with the reference set is informative. In this scenario, it may be necessary to aggregate taxa at a higher level, such as ASVs clustered at 95% similarity or data aggregated to the tribe or genus level. This aggregation would make it more likely that there is some overlap of taxa between the reference and the test data, and thereby give greater differentiation within RPD values.

## Conclusion

The RPD metric provides a single, interpretable measurement of the deviation of a test sample from a specified reference dataset. The advantages of the method include its lack of assumptions about the distribution of the data, its robustness to relative abundance data, its compatibility with downstream statistical analyses, and its basis in the ecological theory that ratios between taxa are informative measurements. In the analyses presented here, RPD performed better than a widely used methodological approach in analyzing four datasets with differing primary outcome variables. Given the applicability of this method to research questions across ecological, environmental, and biomedical science, RPD shows promise as a standardized measurement of compositional variability.

## Supporting information

RPD_SupplementaryMaterials

## Acknowledgements

I received helpful comments on this work from Michael Baym, Trina McMahon, Matthew Stachyra, and Amy Zamora.

## AI Statement

AI was used for the following purposes: creating R code templates for figures; creating R code templates for using functions from the pROC package; proofreading the manuscript and R scripts for inconsistencies; giving a simulated peer review on the manuscript and R scripts.

