## Supplementary material for "Ratio Percentile Deviation (RPD): A nonparametric, compositionally robust method for measuring the divergence of a microbial sample from a reference dataset": RPD_SupplementaryMaterials

### Supplementary Materials for RPD Method

#### Overview:

Several additional analyses were performed to evaluate the robustness of the RPD method. The supplementary materials include the following analyses for the main text dysbiosis results:

- Benchmarking of results against additional predictors
- Robustness of results when selecting a different reference dataset
- Sensitivity analysis changing the reference persistence parameter
- Sensitivity analysis changing the size of the reference dataset
- Estimation of model optimism

All of these analyses use AUC as the outcome of interest. The benchmarking analyses can be directly compared to the analyses in the main text, because they were implemented with the same reference and test datasets. However, while the AUCs in the sensitivity analyses are internally consistent within each analysis, they are not perfectly comparable to the main text results; in these analyses, both the size of the dataset and the fraction of cases in the test dataset was changed to allow repeated random sampling of the datasets.

#### Benchmarking of results against additional predictors

The main text presents RPD compared to BCD for two primary reasons: first, BCD to the reference centroid is a common metric for this type of analysis, and second, BCD to the reference centroid had the best overall performance of the comparator models. For each dataset, I also calculated the AUC for the following predictors (Fig. S1-S3):

- Median BCD of a sample to each reference sample (abbreviated BCDM)
- BCD on the subset of taxa retained in the RPD analysis (BCDS)
- Robust Aitchison distance between each sample and the reference centroid, using the full dataset (AITF)
- Robust Aitchison distance between each sample and the reference centroid, using the subset of taxa retained in the RPD analysis (AIT)
- Euclidean distance on the subset of taxa retained in the RPD analysis (EUC)
- Rank shift analysis using the taxa retained in the RPD analysis (RS)

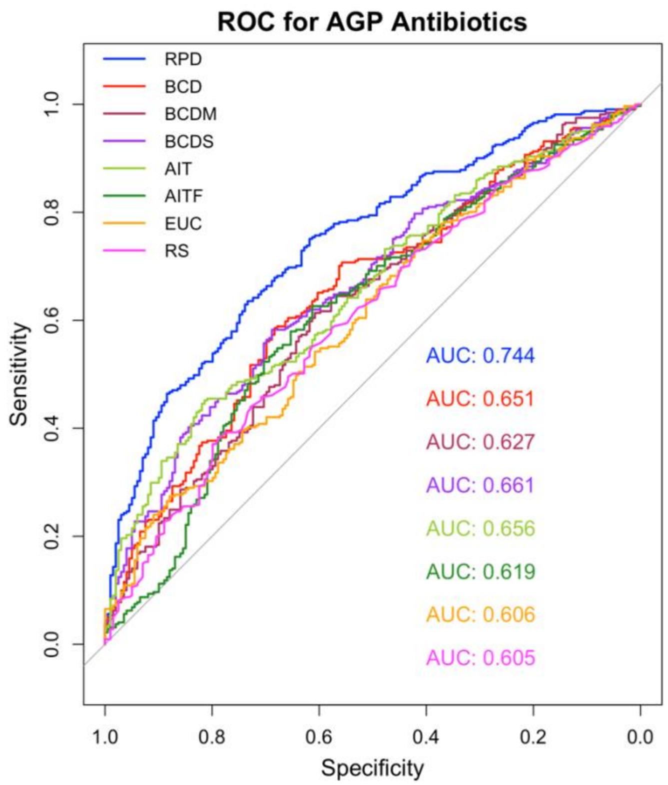

**Figure S1: ROC and associated AUC for AGP Antibiotics with the six additional models**

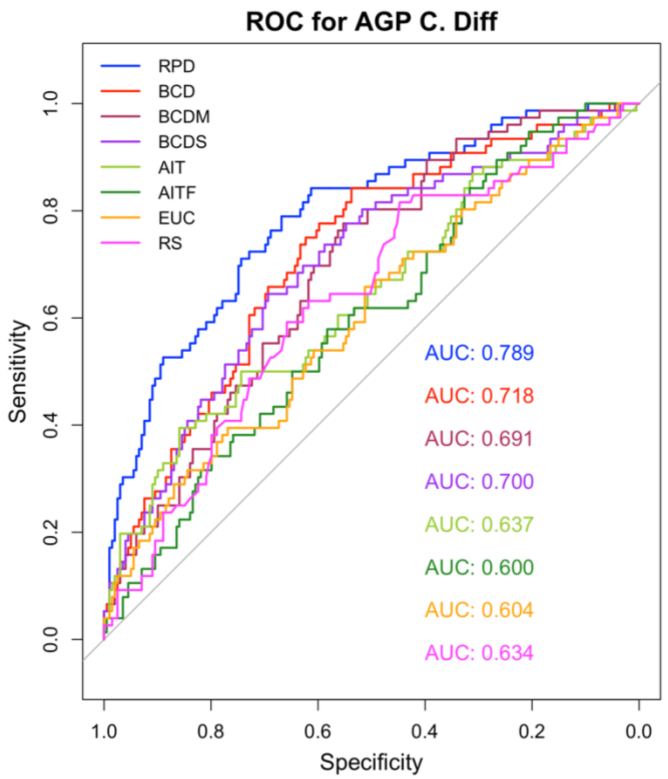

**Figure S2: ROC and associated AUC for AGP *C. difficile* with the six additional models**

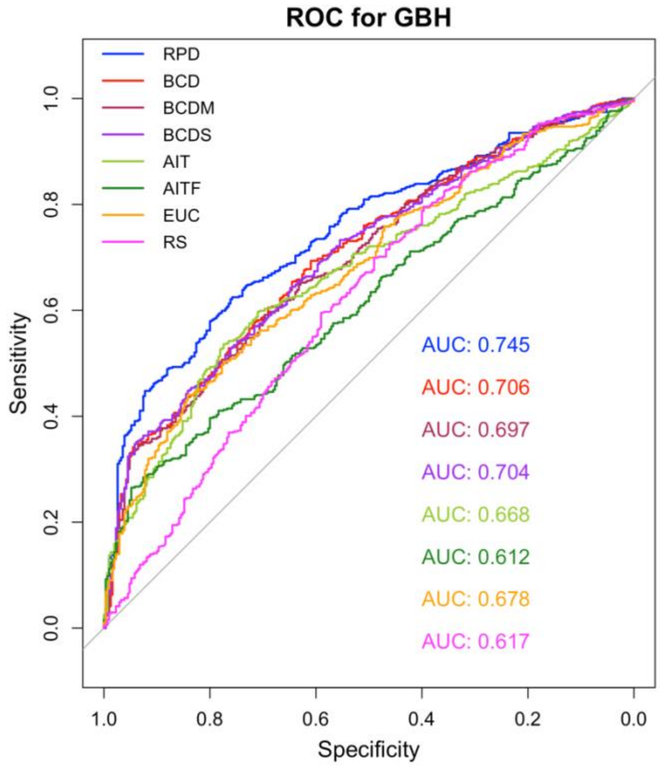


**Figure S3: ROC and associated AUC for GBH with the six additional models**

In all three analyses, RPD as a predictor stands out as better than any benchmark. In two of the three analyses (AGP *C. difficile* and GBH), BCD to the reference centroid was the best-performing comparator model. In the AGP antibiotics dataset, BCD to the centroid calculated on the subset of the dataset was slightly better than BCD to the centroid on the full dataset, but only by 0.01 units. Thus, BCD to centroid on the full dataset had the best overall performance of the benchmarking models. Median BCD to each reference sample was consistently worse than BCD to the reference centroid. Robust Aitchison distance did equivalently well to BCD to the centroid for AGP antibiotics, but performed poorly in the AGP *C. difficile* dataset. Rank shift analysis and Euclidean distance were never competitive predictors.

**<u>Robustness of results when selecting a different reference dataset</u>**

The main text analysis uses 100 random samples of the GBH dataset and 500 random samples of the AGP dataset as reference datasets for the respective analyses. Although the intention with the large sample sizes was to increase the probability of selecting a representative sample, there is still some effect of the specific samples chosen as references.. Here, I repeatedly randomly selected reference datasets of the same size from all healthy individuals. I repeated the analyses presented in the main text 100 times to characterize the distribution of possible results (Fig. S4).

The main outcome of interest was the difference in AUC between the model using RPD as a predictor versus the model using BCD as a predictor. This is a measurement of the superiority of the RPD model. Across the 100 randomizations, the mean improvement in AUC between RPD and BCD was 0.087 for the AGP antibiotics dataset, 0.063 for the AGP *C. difficile* dataset, and 0.030 for the GBH dataset. These improvements in AUC are comparable to the improvements shown in the main text, where the differences were 0.093, 0.071, and 0.039. Furthermore, RPD was a superior predictor to BCD in every iteration; the minimum improvement in AUC was 0.061 for AGP antibiotics, 0.036 for AGP *C. difficile*, and 0.015 for GBH.

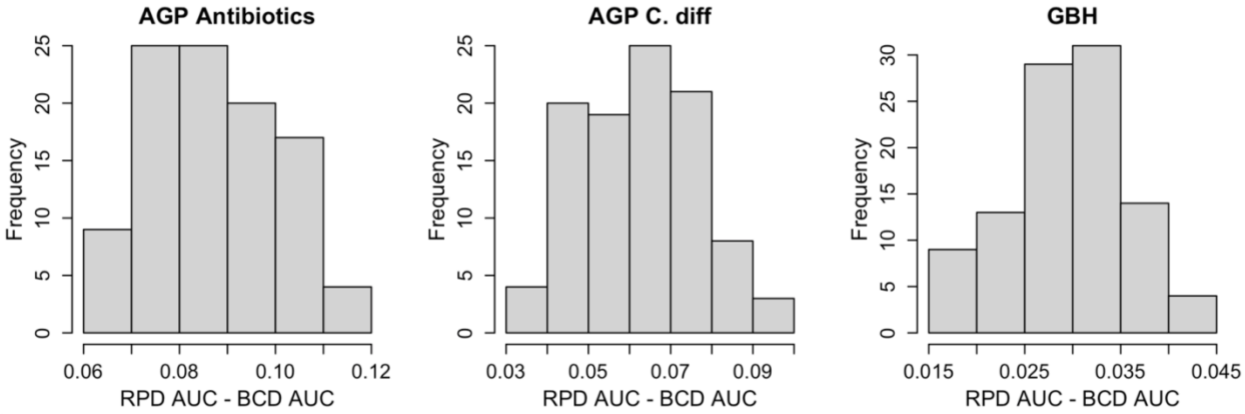


**Figure S4: Difference in AUC between RPD and BCD models when using randomly selected reference samples.** The mean difference in AUC was within 0.01 of the results presented in the main text. RPD was a better predictor than BCD in all cases, seen by all values greater than zero.

**<u>Sensitivity analysis changing the reference persistence parameter</u>**

The reference persistence threshold specifies what fraction of samples a taxon must be present in to be included in the RPD analysis. I first examined what fraction of the community was retained at different reference persistence thresholds. The following analysis changed this parameter in the datasets used in the main text, and calculated the mean and middle 50% of the distribution for the fraction of total abundance retained in the reference communities:

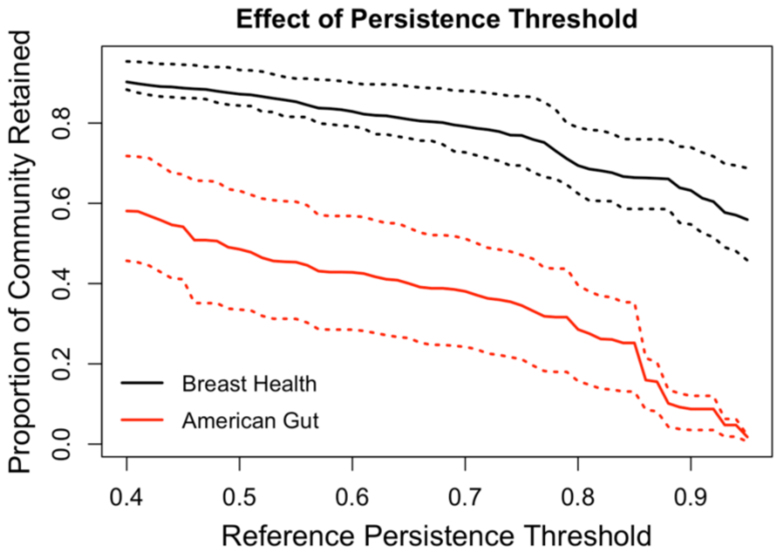

**Figure S5: Fraction of reference communities retained as a function of reference persistence threshold.** Solid lines show the mean fraction of the community retained for the RPD analysis when using difference reference persistence values. Dashed lines show 25th and 75th percentiles.

With the persistence thresholds used in the main text, the analyses retained approximately 40% of the communities for the AGP analyses and approximately 80% of the communities for the GBH analysis. The AGP dataset retained a smaller fraction of the community than the GBH dataset when using the same persistence threshold due to both the greater homogeneity of the GBH dataset and the deeper sequencing of the GBH dataset.

I then systematically changed the reference persistence threshold used in the analyses to evaluate the sensitivity of the analysis results to this parameter. I repeated this analysis with randomly drawn references to characterize the average expected effect across any reference dataset (Fig. S6). For the AGP dataset, I changed the persistence threshold from 0.5 to 0.9. For the GBH dataset, I changed it from 0.6 to 0.95. For this analysis, I used a reference size of 200 for the AGP datasets and 50 for the GBH dataset. The remainder of the healthy subjects were used as the control subjects in the test dataset. Cases were the same as those in the main text. For each reference persistence threshold value, I performed this analysis 50 times for the GBH dataset and 100 times for the AGP dataset.

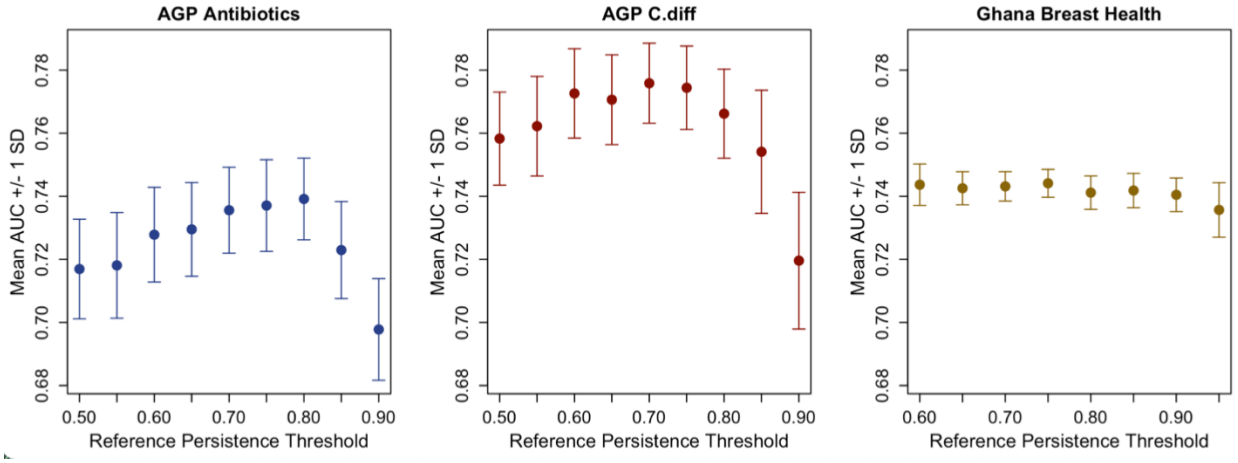


**Figure S6: Effect of the reference persistence threshold on AUC for models using RPD as a predictor.** The AGP datasets show greater sensitivity to the reference persistence parameter, with more variation in AUC over varying persistence thresholds. The optimal reference persistence threshold for the AGP antibiotics dataset is around 0.7 to 0.8, while for the AGP *C. difficile* dataset it is 0.6 to 0.75. For the GBH dataset, values in the range of 0.6 to 0.75 produce similar results, with slightly declining AUC values as higher thresholds. Plots show mean plus/minus one standard deviation.

The AGP datasets were much more sensitive to a change in the reference persistence threshold than the GBH dataset. This is possibly due to the smaller fraction of the total community retained in the AGP dataset. However, the difference in mean AUC values across different persistence thresholds in the AGP analyses was only around 0.02 units when moving across thresholds ranging from 0.5 to 0.8. At higher reference persistence thresholds, model performance suffers.

#### Sensitivity analysis changing the size of the reference dataset

The size of the reference dataset will affect the number of data points in the reference ratio distributions, and therefore will affect the resolution and discrimination of ratio percentiles. To investigate the effect of reference sample size on the main text analyses, I repeated the dysbiosis analyses using smaller reference datasets. Whereas the main text used 500 samples for the AGP analyses and 100 samples for the GBH dataset, I instead used reference sizes of 20, 50, 100, 200, and 400 for the AGP analyses and 20, 50, 100, and 200 for the GBH analysis (400 was not possible due to a smaller control population). The number of control samples was fixed across all analyses to be 200. The cases were equivalent to those used in the main text. I repeated this analysis many times (200 times for the AGP analyses and 100 times for the GBH analysis) for each sample size to characterize the effect of any reference dataset of that size (Fig. S7).

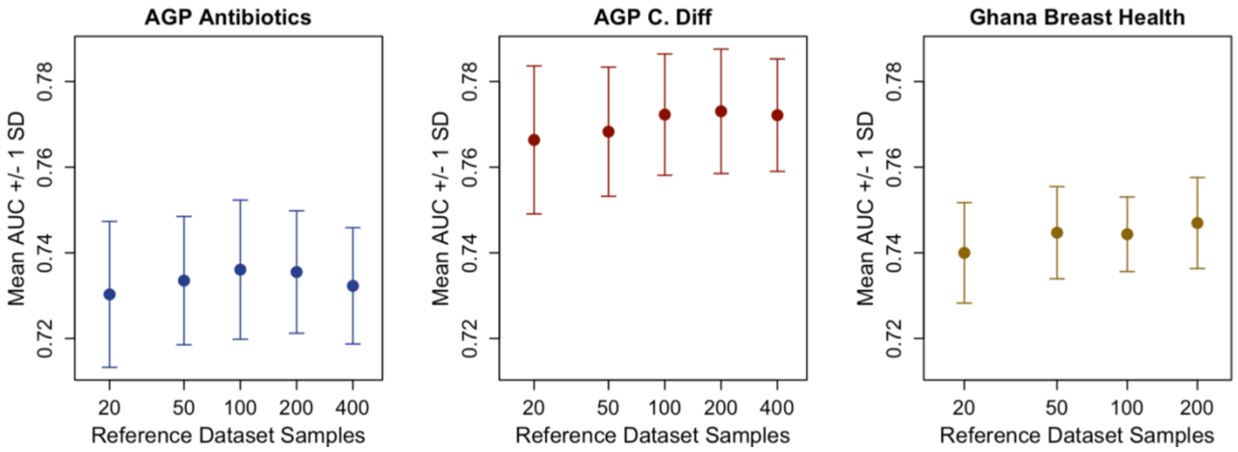


**Figure S7: Effect of reference dataset size on AUC in models using RPD as a predictor.** In the AGP datasets, there was improvement in mean AUC as sample size increased, with most of the benefit realized by the size of 100 samples. In the GBH dataset, there was improvement between 20 and 50 samples, but no meaningful improvement thereafter.

Perhaps surprisingly, the size of the reference dataset had a relatively small effect on AUC, especially when compared to the run-to-run variability or to the effect of the reference persistence threshold. It is possible that this is because 1) sufficiently many taxa are retained in the analyses to still calculate robust averages, or because 2) the case and control samples are sufficiently different that case percentiles are consistently at the very upper or lower extremes. Finally, the slight decrease in mean AUC when moving from a reference size of 200 to a reference size of 400 in the AGP analyses suggests that there may be interplay between the reference persistence threshold and the reference size, where the persistence parameter may need to be adjusted based on reference size.

#### Estimation of model optimism

Model optimism refers to the difference in model performance between training data and unseen data. Here, it is the difference in AUC between models fit on training data versus hold-out data. The main text presents the AUC of the training data, which does not account for model optimism. However, the small number of predictors (only 1) in comparison to the large number of data points suggests that model optimism should be small. To investigate this assumption, I performed a cross-validation of the model on held out data. For 1000 iterations, I divided the test dataset from the main text analysis into two folds of equal size. I trained the model on one fold (containing half the data) and validated it on the other fold by predicting the case vs. control classes from their RPD values. I recorded the AUC of both the RPD and BCD analyses.

The difference in AUC between the training data (main text) and the average of the hold-out AUCs was less than 0.001 in all cases, indicating low model optimism. For the AGP antibiotics dataset, the mean hold-out AUC was 0.7453, less than 0.001 different from the main text analysis. For the AGP *C. difficile* dataset, the mean old-out AUC was 0.7872, a difference of 0.002. And for the GBH dataset, the mean hold-out AUC was 0.7441, a difference of less than 0.001 (Fig. S8).

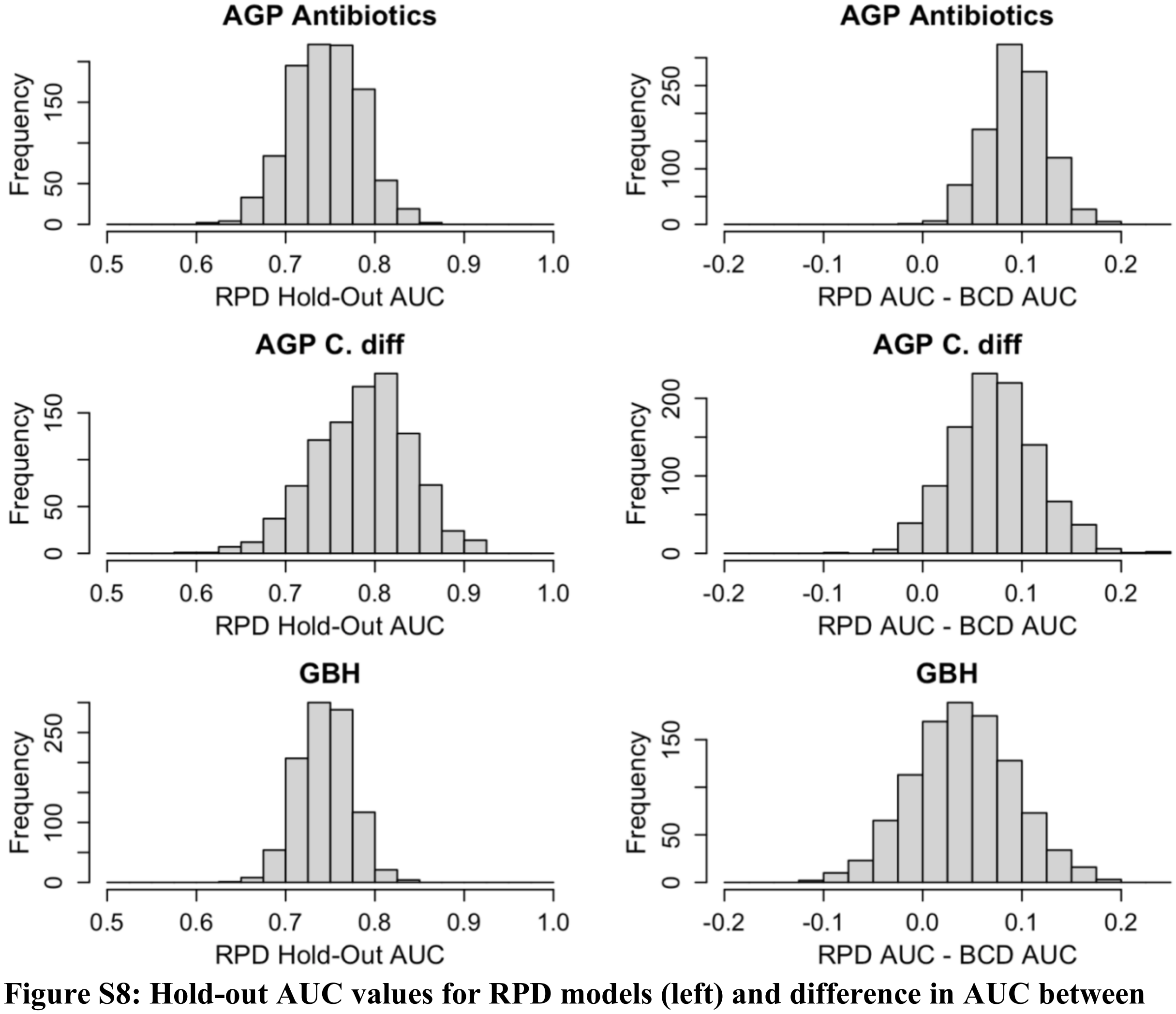


**Figure S8: Hold-out AUC values for RPD models (left) and difference in AUC between hold out RPD models and hold-out BCD models (right).**

Similarly, the average difference in AUCs between the RPD and BCD models applied to held out data were very close to main text values, with mean differences of 0.095 (AGP antibiotics),

0.073 (AGP C. diff), and 0.041 (GBH). However, there was considerably greater variability in the hold-out AUCs. Part of this was due to the application of the model to unseen data, and part of this was due to the smaller sample size used to train the model. Still, RPD was the better predictor across the three datasets, showing a better hold-out AUC in 99.9% of AGP antibiotics runs, 95.5% of AGP *C. difficile* runs, and 78.7% of GBH runs.
